# Signal-independent activation reveals two-component regulatory networks

**DOI:** 10.1101/2024.03.22.586261

**Authors:** Cosme Claverie, Francesco Coppolino, Maria-Vittoria Mazzuoli, Cécile Guyonnet, Elise Jacquemet, Rachel Legendre, Odile Sismeiro, Giuseppe Valerio De Gaetano, Giuseppe Teti, Patrick Trieu-Cuot, Asmaa Tazi, Concetta Beninati, Arnaud Firon

**Author notes:** Correspondence to Arnaud Firon. These authors contributed equally (Cosme Claverie, Francesco Coppolino).

## Abstract

Each bacterial species has specific regulatory systems to control physiology, adaptation, and host interactions. One challenge posed by this diversity is to define the evolving gene regulatory networks. This study aims to characterise two-component systems (TCS) in *Streptococcus agalactiae*, the main cause of neonatal meningitis. Here we demonstrate signal-independent activation of signalling pathways by systematically targeting the conserved mechanism of phosphatase activity of the 14 histidine kinases of the two main TCS families. Transcriptomic analysis resolves most pathways with high resolution, encompassing specialized, connected, and global regulatory systems. The activated network notably reveals the connection between CovRS and SaeRS signaling through the adhesin PbsP, linking the main regulators of host interactions to balance pathogenicity. Additionally, constitutive activation of the BceRS system reveals its role in cell envelope homeostasis beyond antimicrobial resistance. Overall, this study demonstrates the generalizability and versatility of TCS genetic activation to uncover regulatory logics and biological processes.

## INTRODUCTION

Two-component systems (TCSs) are one of the main bacterial signaling mechanisms. In their simplest form, an environmental signal activates a histidine kinase (HK), which phosphorylates a cognate response regulator (RR), leading to the transcription of specific genes that mediate the cellular response to the stimuli ^1^. Actually, TCSs are sophisticated molecular machineries with buffering and insulating mechanisms that dynamically control specific or global cellular responses ^2–4^. Considerable effort has been made to define TCS regulatory networks in both model and pathogenic species, including by comprehensive analysis ^5,6^. Although knowledge gained in one species can provide information about homologous systems, TCS are characterized by their diversity, plasticity and evolvability ^7,8^. This prevents global inferences even between closely related species ^9,10^. This evolution of regulatory networks is sustained by several mechanisms, including mutations, horizontal gene transfer, duplication follow by neofunctionalization, and rewiring that shapes adaptation and speciation ^11–13^.

Functional, evolutionary, and system analyses require characterizing individual signaling pathways and integrating them into the cellular regulatory network. Traditionally, regulons are characterized using inactivated TCS mutants. One common pitfall is that TCSs are not active until the specific, but usually unknown, stimulus is provided. Current approaches to overcome signal requirement are based on phosphomimetic mutation of the RR ^14,15^ and profiling of protein-DNA interaction ^16–18^. An alternative approach exploits the distinct HK enzymatic activities. The HK cytoplasmic core, called the transmitter module, is composed of the DHp (Dimerization and Histidine phosphotransfer) and CA (catalytic and ATP-binding) domains ^19–21^. The two domains are dynamically structured in specific conformations that catalyse three distinct reactions: autophosphorylation of a conserved histidine residue in the DHp domain, phosphotransfer to a conserved aspartate on the RR, and RR dephosphorylation.

Pioneering studies have identified mutations abolishing the HK phosphatase activity leading to increased RR phosphorylation and signaling pathway activation ^22–24^.

The importance of HK phosphatase activity *in vivo* has been initially debated, especially when considering the lability of RR phosphorylation and spontaneous dephosphorylation rate _25_. Nowadays, the phosphatase activity is recognized as essential for the dynamics of the response and to ensure that the RR is activated by the cognate HK only ^26,27^. Co-evolving residues and HKs conformational rearrangements ensure specificity and directionality of enzymatic reactions ^28–30^. While the activation mechanism involving the conserved histidine residue is fundamentally conserved among HKs, the phosphatase mechanism has remained more elusive due to variations in the DHp domain ^31^. Then, a seminal study has proposed a conserved phosphatase mechanism for the two main HisKA and HisKA_3 family, identifying conserved motifs and specific catalytic residues needed for the correct positioning of nucleophilic attack ^31,32^. Substitution of the catalytic residues abolishes the phosphatase activity without impacting the autokinase and phosphotransfer activities, resulting in increased RR phosphorylation and pathway activation for the individual HKs reported to date ^32–37^.

This study aims to systematically test the proposed conserved mechanism of phosphatase activity, the *in vivo* effect of phosphatase-deficient HK, and the activation of the regulatory network in all HisKA and HisKA_3 systems in a bacterium. We focused on *Streptococcus agalactiae* (Group B *Streptococcus*, GBS), a pathobiont that is commensal in adults but pathogenic during pregnancy and in neonates, for whom it is the leading cause of invasive infections ^38,39^. We report that targeting HK phosphatase activity provides high-resolution views of signaling pathways for most TCSs independently from environmental signals. In addition, regulatory network activation resolves the connectivity between TCSs involved in host-pathogen interactions and reveals the physiological function of a TCS involved in antimicrobial resistance. This systematic analysis argues for the widespread adoption of this gain-of-function approach to decipher TCSs signaling in genetically manipulable species.

## RESULTS

### The HK^+^ collection targets the phosphatase activity of Histidine Kinase

We undertook a genetic approach to systematically test the hypothesis of a conserved dephosphorylation mechanism in the two major HK families ^32^. The genome of the BM110 strain belonging to the hypervirulent clonal complex 17 (CC-17) encodes 20 HKs ^40^, among which 12 and 2 have a HisKA and HisKA_3 DHp domain, respectively (Supplementary Table 1). Their H-box motif always contains the conserved phospho-acceptor histidine, immediately followed by the predicted phosphatase motif (Fig. 1A). Eleven of the twelve HisKA proteins have the E/DxxT/N motif with a putative threonine catalytic residue, while the remaining HisKA protein (BceS) has a divergent sequence composition (QMKV) with a valine at the predicted catalytic position (Fig. 1A). The two HisKA_3 proteins have the DxxxQ/H motif with the predicted glutamine or histidine catalytic residue (Fig. 1A). The 14 HKs encoding genes are organized in operon with their cognate response regulator (RR) belonging to the OmpR (with His_KA) or LuxR (with His_KA3) family, but one system is not functional (HK10655- RR10650^fs^) due to a pseudogenization of the RR in the CC-17 hypervirulent GBS lineage (Supplementary Table 1).

**Figure 1:**
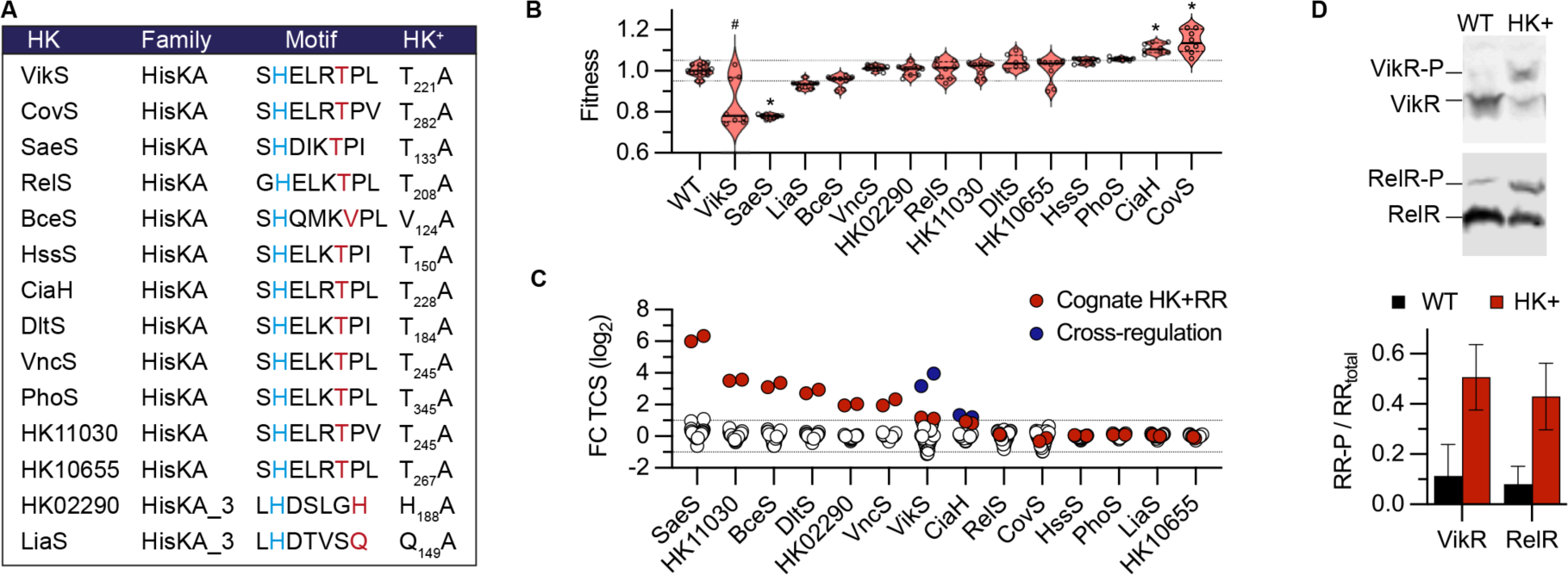
Mutation of HK phosphatase catalytic residue activates TCS signaling. **A.** Conserved motif of the HisKA and HisKA_3 histidine kinases with the phospho-acceptor histidine (blue) and the predicted residue specifically involved in the phosphatase activity (red). The phosphatase residue is substituted by an alanine in the HK^+^ mutants. **B.** Fitness of HK^+^ mutants. The violin plots represent the distribution of the relative doubling time (F = D_x_ / D_WT mean_) in exponentially growth phase in rich medium with the median (bar) and the interquartile range (dashes). Data are from biological replicate (n = 16 for the WT, n = 8 for mutants) and significant differences are highlighted (*, |F | > 0.1, Mann Whitney test p < 10^-4^). The bimodal distribution due to the occurrence of faster-growing VikS_T221A_ suppressors is highlighted (#). **C.** Activation of transcriptional feedback loops in HK^+^ mutants. Fold changes (FC) for all genes encoding TCS (n = 41) in each HK^+^ mutant after RNA-seq analysis are shown as dots. The HK- RR gene pair in the corresponding HK^+^ mutant is highlighted in red (*e.g.*, *saeRS* in SaeS_T133A_). Cross-regulations, defined as significant differential expression of a TCS gene pair not corresponding to the HK^+^ mutation, are highlighted in blue (*hk11050*-*rr11055* in VikST_221A_ and *relRS* in CiaH_T228A_). **D.** Activation of the VikR and the RelR response regulators by phosphorylation in the corresponding HK^+^ mutants. Upper: representative Phos-Tag western-blots with anti-FLAG antibodies allowing to separate phosphorylated and non-phosphorylated forms of the ectopically expressed epitope-tagged RR. Bottom: quantification of the proportion of phosphorylated regulator in the WT (black) and the cognate HK^+^ mutant (red). Bars represent the mean with SD of biological replicate (n = 3).

We generated 14 strains, called the HK^+^ collection, with an alanine substitution of the predicted phosphatase catalytic residue (Fig. 1A). Whole-genome sequencing confirmed the chromosomal substitution of targeted base pairs and absence of secondary mutations in 11 out of the 14 HK^+^ strains. In the three remaining HK^+^ (CovS_T282A_, VikS_T221A_, and RelS_T208A_), we sequenced independent mutants and selected one with a single secondary mutation (Supplementary Table 2). Noteworthy, the selected VikS_T221A_ mutant has a non-synonymous polymorphism in the glutamine transporter GlnPQ that we cannot exclude as a compensatory mutation. Four independent VikS_T221A_ mutants also have putative compensatory mutations (Supplementary Table S2), as frequently observed in mutants in the homologous WalRK system essential for cell wall remodelling during growth and division ^41–43^.

Individual growth curves show a significant effect (|F| > 0.1, Mann Whitney test p < 10^-4^) of the HK^+^ mutation for four mutants, two gaining a reproductible fitness advantage (CovS_T282A_ and CiaH_T228A_) and two having specific phenotypes (Fig. 1B). Not surprisingly, the slow growing VikS_T221A_ mutant is unstable and gives rise to faster-growing cultures likely due to additional mutations (Supplementary Fig. S1). In contrast, the SaeS_T133A_ mutant exhibits a density-dependent phenotype characterised by a decreasing growth rate in the exponential phase and a lower final OD (Supplementary Fig. S1). Additionally, two mutants have increased antibiotic susceptibilities: the VikS_T221A_ mutant against beta-lactams, in agreement with a conserved function in cell wall metabolism, and the RelS mutant against fosfomycin (Supplementary Table S3).

### HK^+^ activate positive feedback loops

To test TCS activation, we first relied on positive feedback loops. This autoregulation is often observed through direct transcriptional activation of the TCS operon by the activated RR ^2^. We therefore analysed the transcription of all HKs and RRs encoding genes (n = 41, including non-His_KA and His_KA3 TCSs and an orphan RR) in each HK^+^ mutants by RNA-sequencing from cultures grown in a standardized condition (THY, 37°C, OD_600_ = 0.5). A positive feedback loop, defined by a FC > 2 and p-adj < 10^-4^ for the HK and RR genes, is observed in seven HK^+^ mutants (Fig. 1C). Furthermore, two TCSs are significantly regulated in an unrelated HK^+^ mutant: the HK11050-RR11055 system, which does not contain a His_KA and His_KA3 domain, in the VikS_T221A_ mutant and the RelRS system, which is not positively auto-regulated, in the CiaH_T228A_ mutant (Fig. 1C). As an independent approach to test TCS activation, we introduced in each mutant a vector expressing an epitope-tagged copy of the cognate regulator. For two mutants (VikS_T221A_ and RelS_T208A_), an increased level of phosphorylation of the ectopically expressed regulator is detected in the HK^+^ mutant compared to the WT strain after Phos-Tag electrophoresis and western analysis with anti-FLAG antibodies (Fig. 1D). Overall, by considering epitope-tagged RR activation by phosphorylation and positive feedback loops, the majority (8/14) of HK^+^ mutations appear to activate the corresponding TCS signaling pathway.

### The activated gene regulatory network.

To characterise the activated pathways, we analysed the RNA-seq profiles of each HK^+^ mutant grown under standardised conditions, independent of specific environmental cues (*i.e.,* exponential phase in rich media at 37°C). As illustrated with HK11030_T245A_ and VncS_T245A_ (Fig. 2A), six HK^+^ mutants (including SaeS_T133A_, BceS_V124A_, HK02290_H188A_, and DltS_T184A_) are associated with highly significant activation of specific genes (p-adj < 10^-250^) (Supplementary Fig. S2). Four additional mutants (RelS_T208A_, CiaH_T228A_, VikS_T221A_, and CovS_T282A_) are associated with an intermediate activation of larger gene sets (p-adj > 10^-150^), while the remaining four (HssS_T150A_, LiaS_Q149A_, PhoS_T345A_, and the HK10655_T267A_ with a frameshifted RR) gave no or low significant signals (p-adj > 10_-10_) (Supplementary Fig. S2).

**Figure 2:**
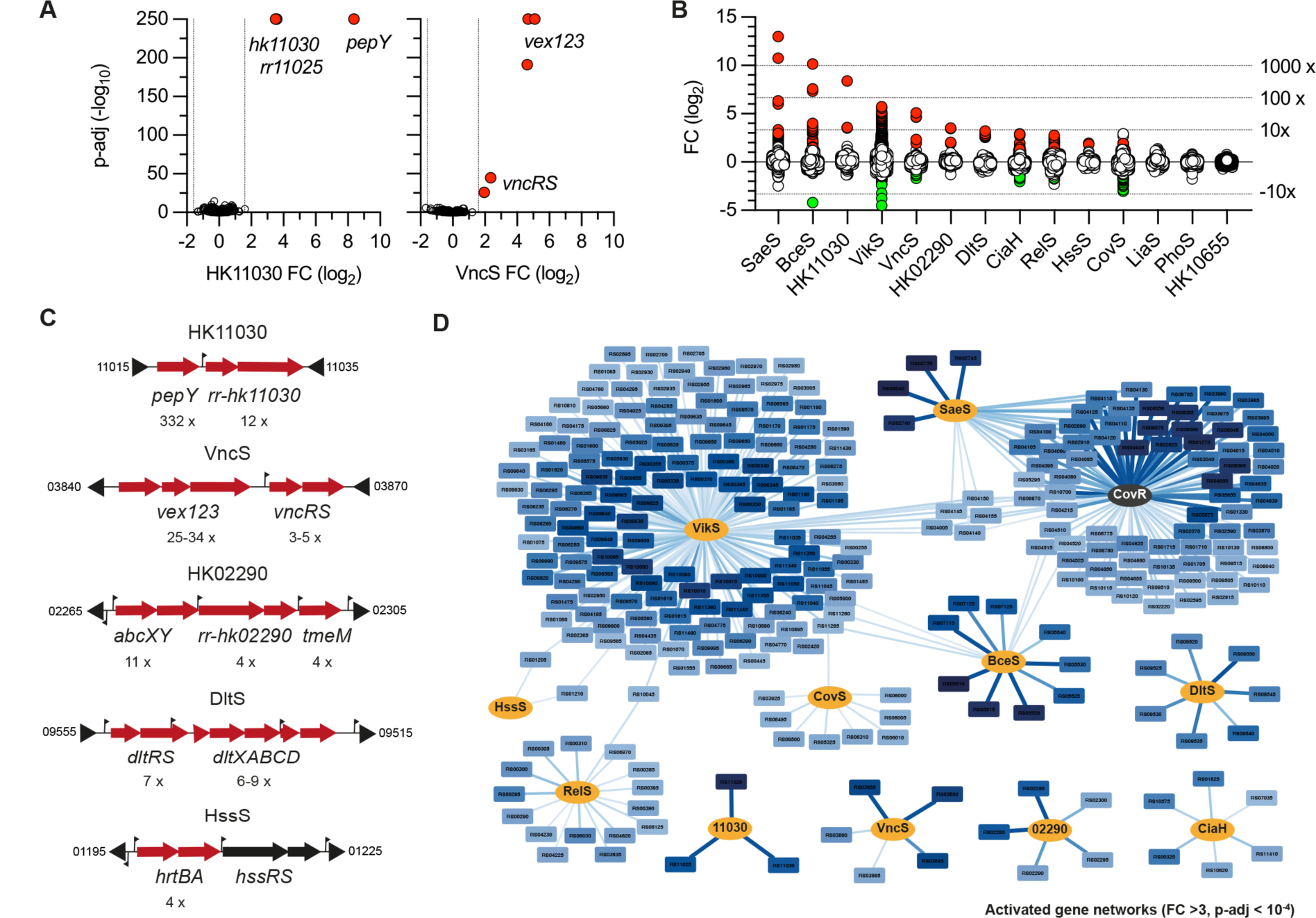
The activated gene regulatory networks. A. Volcano plot of significant differential gene expression in the HK11030_T245A_ (right panel) and VcnS_T245A_ (left panel) mutants. Transcriptomes by RNA-seq against the WT strain were done in exponential growth phase in rich media (THY). Red dots highlight significantly differentially regulated genes above the thresholds FC > 3, p-adj < 10^-4^. Volcano plots for the 14 HK^+^ mutant are provided in the related Supplementary Figure S2. B. Violin distribution of transcriptional fold change in the 14 HK^+^ mutants. Coloured dots represent significantly activated (red) and repressed (green) genes (|FC| > 3, p-adj < 10^-4^), respectively. C. Activated chromosomal loci in selected HK^+^ mutants. Fold changes are indicated below the activated genes (red arrows). Transcriptional start sites identified by genome wide TSS mapping are represented by vertical flags. NCBI gene ID bordering the loci are shown in a shortened form (e.g.: 11015 = BQ8897_RS11015). Activated loci for each HK^+^ mutant are provided in the related Supplementary Figure S3. D. Network of activated genes. Histidine kinases (orange nodes) are connected to their activated genes (light to dark blue nodes). Edge thickness and gene node colour are proportional to statistical significance and fold change, respectively. Activated genes in the CovR_D53A_ mutant (black node) are included to account for the specificity of CovR as a global repressor.

Since most RRs are transcriptional activators, we focused the analysis on activated genes. By applying strict thresholds (FC > 3, p-adj < 10^-4^) for normalisation between samples and excluding genes with very low read counts in all samples and genes localised in mobile genetic elements, 219 genes (11.9% of the 1838 genes analysed) are transcriptionally activated in at least one HK^+^ mutant (Supplementary Table S4E). Transcriptional activation can be up to 8000-fold, with an average fold change of 61.6-fold and an uneven distribution between HK^+^ mutants (Fig. 2B).

The number of activated genes ranges from 3 (HK11030_T245A_) to 139 (VikS_T221A_) (Fig. 2B and Supplementary Table S4). Five regulatory systems activate a specific genetic program, four of them (HK11030_T245A_, HK02290_H188A_, VncS_T245A_, DltS_T184A_) positively regulating a single functional genetic module composed of their own operon and at least one additional gene involved in the cellular response localized into, or adjacent to, the TCS operon (Fig. 2C), and one system (CiaH_T228A_) coordinating the activation of at least six independent loci (Supplementary Fig. S3). Four additional TCSs activate specific genes but share 1 to 3 activated genes with the VikS_T221A_ mutant (Fig. 2D). One of these connected systems (HssS_T150A_) is specialised in haem detoxification via the transcriptional activation of the *hrtBA* genes encoding a specific ABC transporter ^44^, which is similarly activated in the VikS_T221A_ mutant. The three additional connected systems activate several loci involved in host-pathogen interaction (SaeS_T133A_: adhesins and secreted proteins), drug resistance (BceS_V124A_: transporters and peptidase), or nucleotide metabolism (RelS_T208A_: *de novo* purine synthesis and ectonucleotidases), with (SaeS_T133A_, BceS_V124A_) or without (RelS_T208A_) a positive feedback loop (Fig. 2D and Supplementary Fig. S3).

### Positive and negative interaction between TCS systems.

Overall, the HK^+^ mutation activates the signaling pathway for 10 out of 12 TCSs (Fig. 2E), excluding the CovS_T282A_ repressing system analysed separately and the negative control HK10655_T267A_ with a frameshifted RR. Notably, each HK^+^ mutant is associated with the activation of specific genes, except the global regulator VikRS (Fig. 2E and Supplementary Table S4E). As expected, the VikS_T221A_ regulon included several operons involved in cell wall metabolism (Supplementary Fig S3). However, constitutive activation of VikS probably leads to the activation of related stress and cell-wall signaling pathways. To identify relationships between TCS pathways involved in related processes, we analysed the 219 genes activated in at least one HK^+^ mutant for their expression in the whole RNA-seq dataset. This analysis confirmed the partial activation of SaeRS signaling in the VikS_T221A_ mutant (a shared CovR connection in fact, see specific section below) sustained by genes with 1<FC<3 and significant but higher p-adj value compared to the SaeS_T133A_ activated system (Supplementary Table S4). Similarly, by considering significantly regulated genes with lower thresholds (1<IFCI< 3, p- adj<0.05), significant positive or antagonistic interactions were detected between signaling pathways (*e.g*., DltS activating CiaH and VikS, CiaH antagonising RelS, HK02290 antagonising HK11030). Finally, relaxing the thresholds also reveals the first five genes of the *phoRS* operon as the most and only significantly up-regulated genes (1,5<FC<1,75; 7.10^-3^< p- adj<10^-5^) in the PhoS_T345A_ mutant (Supplementary Table S4D), suggesting a conserved mechanism of phosphatase activity but an inefficient activation of the PhoR regulator in the corresponding HK^+^ mutant.

### Activation of the global repressor of virulence CovRS.

The CovRS system plays a critical role in regulating GBS virulence by acting as a global repressor ^45^. Analysis of RNA-seq to identify negative regulation using similar thresholds (- 3>FC, p-adj < 10^-4^) revealed the repression of 32 genes, primarily in VikS_T221A_ (17 genes) and CovS_T282A_ (14 genes) mutants (Supplementary Table S4). Notably, the most highly repressed gene (22-fold by VikRS) encodes a D-L endopeptidase, reminiscent of WalRK homologous systems ^41^. However, only 5 of the repressed genes in the CovS_T282A_ mutant belong to the CovR regulon previously characterized with inactivated mutants and genome-wide binding experiments ^45^. Comparative analysis between CovS_T282A_ (activated) and CovR_D53A_ (inactivated) transcriptomes shows the activation of CovRS signaling when the CovR repressor is inactive without significant over-repression in the HK^+^ mutant (Fig. 3A). Nevertheless, a significant inverse correlation between the CovS_T282A_ and CovR_D53A_ transcriptomes is observed for genes that do not belong to the direct CovR regulon (Fig. 3B), suggesting that overactivation of CovR primarily increases binding to co-regulated promoters and low-affinity binding sites, and in agreement with CovR being already significantly phosphorylated in the WT strain ^45^.

**Figure 3:**
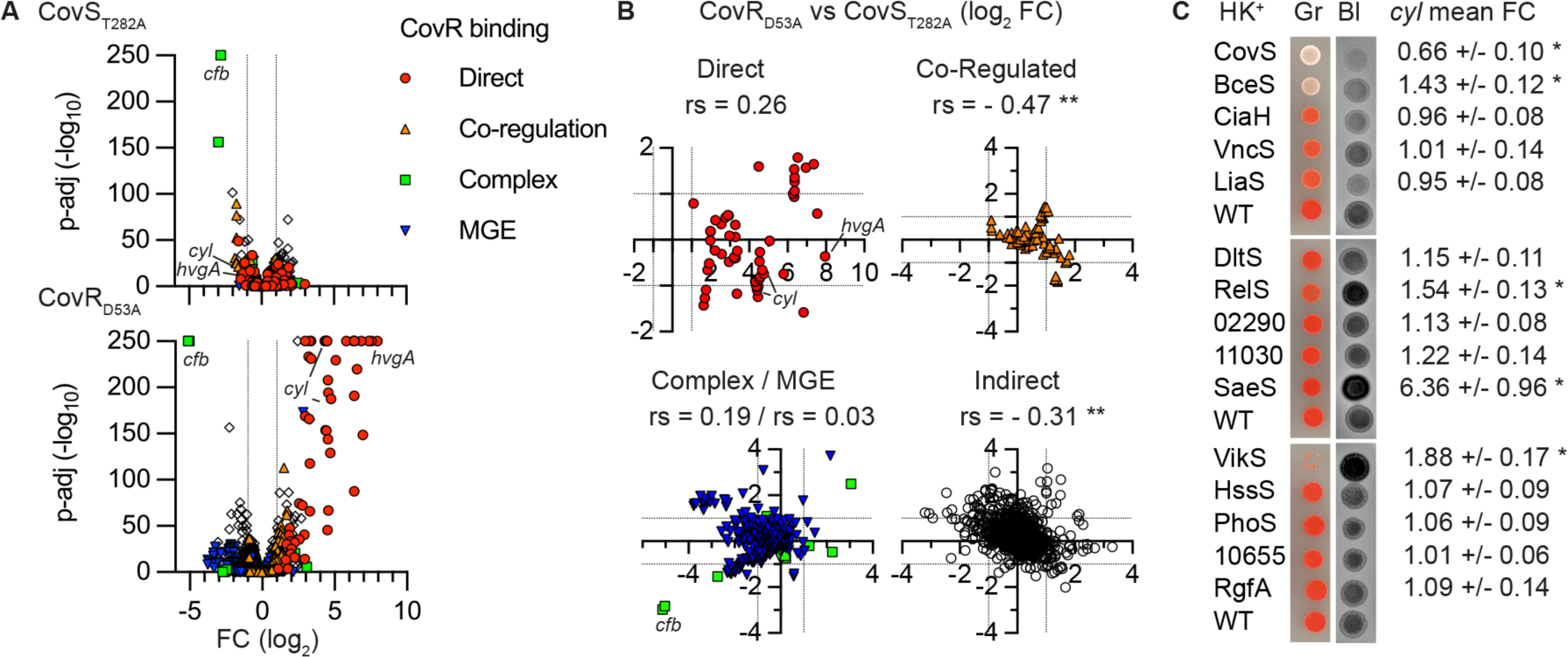
Activation of the global repressor of virulence CovR A. Volcano plot of significant fold changes in the CovR-active (CovS_T282A_) and CovR-inactive (CovR_D53A_) mutants. Coloured dots highlight genes according to CovR-regulatory mechanisms as previously defined by genome-wide binding: direct repression (red), CovR-binding requiring additional regulators for activation (orange), atypical CovR-binding inside ORFs or positive regulation (green), silencing or anti-silencing of genes in mobile genetic elements (blue). B. Comparison of fold changes between CovR inactivation and activation according to regulatory mechanisms. Significant correlations are highlighted (**: rs non-parametric spearman correlation, p (one-tailed) < 10^-4^). C. Pigmentation and haemolytic phenotypes of the HK^+^ mutants on selective media. Spots of diluted cultures are incubated on Granada (Gr) and Columbia horse blood (Bl) plates in anaerobiosis and aerobiosis, respectively. Haemolytic activity is visualized by the dark halo on the inverted black and white photographs. The BM110 parental strain (WT) was added on each plate as control. Note that the VikS_T221A_ mutant does not grow on Granada media, the basis of this phenotype requiring further investigation. The mean RNA-seq fold change with SD of the 12 genes *cyl* operon encoding the pigmented haemolysin ß-h/c directly repressed by CovR are indicated (* p-adj < 0.005).

The CovS_T282A_ mutation offers insight into the complexity of the CovR regulatory network but is less informative than the inactivation of the system (Fig. 3A). We therefore included the CovR_D53A_ transcriptome in the HK^+^ dataset. The global gene network reveals connections between CovR-repressed genes and SaeS_T133A_, VikS_T221A_, or BceS_V124A_ activated- genes (Fig. 2D). Notably, SaeS_T133A_ is highly connected with the direct CovR-repressed genes, while BceS_V124A_ activates only three CovR indirectly regulated genes. Functional assays using pigmented beta-haemoly sin/cytolysin (ß-h/c) production as a natural reporter of CovR activity first confirmed the non-pigmented and non-haemolytic phenotypes of the CovS_T282A_ mutant (Fig. 3C), in agreement with CovR directly repressing the *cyl* operon encoding the ß-h/c synthesis and export machineries ^45–47^. The phenotype of six additional HK^+^ mutants are different from the WT strain on selective media, either increasing (SaeS_T133A_, VikS_T221A_, and RelS_T208A_) or decreasing (BceS_V124A_, CiaH_T228A,_ and LiaS_Q149A_) pigmentation and/or haemolytic activity (Fig. 3C). However, the lack of correlation between the transcription of the *cyl* operon and the pigmentation/haemolytic phenotypes in several HK^+^ mutants (Fig. 3C) suggests additional post-transcriptional regulatory mechanisms influencing ß-h/c activity, particularly in mutants with altered cell surface composition in which the toxin interacts ^48^.

### The PbsP adhesin connects SaeRS and CovRS signaling.

We sought to decipher the connection between the CovRS and SaeRS systems, two main regulators of host-pathogen interactions. Published transcriptomes with *saeRS* deletion mutants define a large regulon of 400-600 genes depending on growth conditions ^49^. In contrast, analysis of the SaeS_T133A_ transcriptome reveals the highly significant (63 < FC < 8080-fold, p-adj < 10^-^ _250_) activation of four genes only, along with a partial activation of the CovR regulon (Fig. 4A). We confirmed the stratification of the SaeS_T133A_ differentially regulated genes by RT-qPCR, validating 3 groups: the *pbsP* and *bvaP* genes, the *saeRS* operon, and the CovR-regulated genes represented by the directly repressed genes *cylE* and *hvgA* (Fig. 4B).

**Figure 4:**
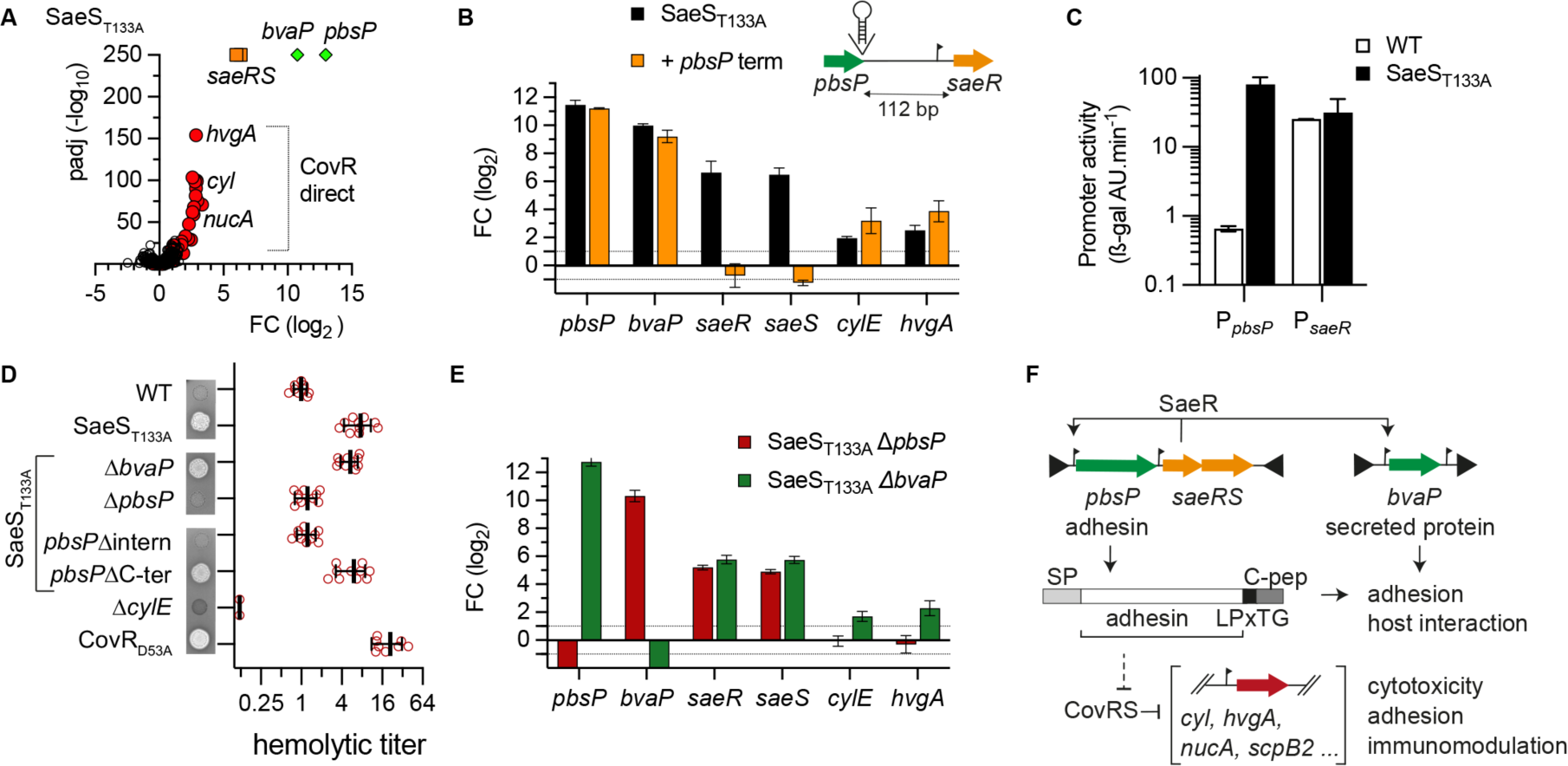
Adhesin-dependent wiring of the SaeR and CovR regulatory networks. A. Volcano plot of significant fold changes in the SaeS_T133A_ mutant. Dot colours highlight the stratification of activated genes between *pbsP* and *bvaP* (green), the *saeRS* operon (orange) and the CovR-regulated genes (red). B. Indirect positive feedback loop of the *saeRS* operon. The *pbsP* and *saeRS* genes are separated by a 112 bp intergenic region containing a *saeR* transcriptional start site located 31 bp from the SaeR start codon. Transcriptional activation of *pbsP* and *saeRS* is uncoupled by the integration of a canonical promoter at the 3’ end of *pbsP*. Fold change of selected genes are quantified by RT-qPCR in the SaeS_T133A_ (black) and in the SaeS_T133A_ + *pbsP* terminator (orange) mutants. Mean and SD are shown for biological replicate (n = 3). C. Activities of the P*_pbsP_* and P*_saeR_* promoters in the WT and SaeS_T133A_ mutant. Bars represent the activity of the ectopic ß-galactosidase reporter system under the control of the tested promoters in the WT and the SaeS_T133A_ mutant. Mean and SD are shown for biological duplicate (n = 2). D. Hyper-haemolytic activity of the SaeS_T133A_ mutant is dependent on the PbsP adhesin. Qualitative and semi-quantitative haemolytic activity are tested on Columbia blood agar media and with defibrinated horse blood, respectively. The Δ*cylE* and CovR_D53A_ mutants are included as negative and positive controls, respectively. Haemolytic titres are normalized against the WT strain. Independent experiments are shown by dots with mean and SD for the biological replicate (n = 8). E. Upregulation of the PbsP adhesin activates CovR-regulated genes. Transcriptional fold change of selected genes by RT-qPCR in the SaeS_T133A_ Δ*pbsP* (red) and SaeS_T133A_ Δ*bvaP* (green) double mutants. Mean and SD are shown for biological replicate (n = 3). F. Wiring diagram of the SaeRS signaling pathway. The *saeRS* operon (orange) is transcribed at a basal level by a constitutive promoter. Upon TCS activation, the SaeR regulator activates the transcription of genes encoding the PbsP and BvaP virulence factors (green), and indirectly its own operon through a *pbsP* terminator readthrough. The over-expression of the PbsP adhesin domain, but not the carboxy-terminal part containing the LPxTG anchoring motif and the hydrophobic C-peptide, is necessary to trigger CovR-regulated virulence factor expression.

Intrigued by the almost 50-fold difference between *pbsP* and *saeRS* up-regulation, we analysed the genomic locus in detail. The 112 bp *pbsP*-*saeRS* intergenic region contains a P*_saeR_* promoter but no canonical transcriptional terminator. The integration of such a terminator precisely after the *pbsP* stop codon in the SaeS_T133A_ mutant abolishes *saeRS* overexpression while having no impact on other activated genes (Fig. 4B). Quantification of promoter activities using *ß-*galactosidase reporters confirms a similar activity of P*_saeR_* in the WT and in SaeS_T133A_ mutant, and the strong activation of P*_pbsP_* upon activation of the SaeRS system (Fig. 4C). This shows an indirect positive feedback loop of the *saeRS* operon, which is transcribed by its constitutive promoter and regulated by *pbsP* termination readthrough. Interestingly, the basal level of *saeRS* transcription in SaeS_T133A_ with the *pbsP* terminator is sufficient to fully activate *pbsP* and *bvaP* (Fig. 4B), implying that the indirect feedback loop may be physiologically relevant only to control the dynamic, and not the amplitude, of the response.

We next analysed the connection between SaeRS and CovRS signaling. The activation of CovR-regulated genes in the SaeS_T133A_ mutant is intermediate when compared to the CovR_D53A_ mutant (Fig. 3A and Fig. 4A). One hypothesis could be a competitive binding between SaeR and CovR, but it is unlikely that all binding sites will allow both SaeR-activation and CovR-repression. As an alternative, we hypothesized that the two genes specifically regulated in the SaeS_T133A_ mutant, encoding the PbsP cell-wall anchored adhesin ^50,51^ and the BvaP secreted protein ^52^, could be involved in the activation of the CovR regulon. Indeed, deletion of *pbsP*, but not of *bvaP*, in the SaeS_T133A_ mutant restores a WT haemolytic activity (Fig. 4D). In agreement with the phenotypes, the deletion of *pbsP* in the SaeS_T133A_ mutant restores a WT level of the CovR-regulated genes *cylE* and *hvgA*, while the *saeRS* and *bvaP* genes are still similarly up-regulated (Fig. 4E).

After cleavage by the enzyme sortase A and anchoring to the cell wall, the remaining carboxy-terminal domain of an LPxTG adhesin can acts as a signaling molecule by interacting with the transmembrane domain of a specific HK ^53^. We therefore considered this C-peptide mechanism and constructed mutants expressing truncated PbsP variants in the SaeS_T133A_ mutant. In-frame deletion of the PbsP C-peptide (*e.g.,* 108 bp deletion including the LPxTG cell-wall anchoring motif until the penultimate codon) has no effect on the induction of the CovR-regulated haemolytic activity (Fig. 4D). In contrast, in-frame deletion of the PbsP adhesin domain (1239 bp deletion leaving the signal peptide and the LPxTG cell wall anchoring motif intact) restores the haemolytic activity of the SaeS_T133A_ mutant to WT level (Fig. 4D). Furthermore, the growth defect of the SaeS_T133A_ mutant, which is similar to the growth defect of the CovR_D53A_ mutant, is suppressed by deletion of *pbsP* or of the adhesin part of *pbsP* (Supplementary Fig. S4). Thus, the PbsP adhesin domain triggers CovR signaling either by interacting with CovS or co-regulatory proteins ^47^ or by inducing surface perturbations specifically sensed by the CovRS system (Fig. 4F).

### Drug-independent activity of the BceRS three-component system.

We next sought to decipher the function of the BceRS system which belongs to a conserved TCS family that relies on a transporter to sense and transmit environmental signals to the HK ^54^. Transcriptome of the BceS_V124A_ mutant reveals a 9-gene regulon, including the *bceRS* operon and adjacent genes (Fig. 5A and Supplementary Fig. S2 and S3). Further validation by RT-qPCR confirmed the 10 to 1000-fold activation of the regulon in absence of drugs in the BceS_V124A_ mutant (Fig. 5B). As expected, mutation of the BceR regulator to a non-phosphorylated form (BceR_D55A_) abolishes the activation of the signaling pathway in the BceS_V124A_ mutant (Fig. 5B). Interestingly, deletion of the transporter/sensor (Δ*bceAB*) in the BceS_V124A_ mutant also switches off the signaling pathway (Fig. 5B), showing the essential role of the BceAB transporter in activating BceRS signaling in absence of inducing signals.

**Figure 5:**
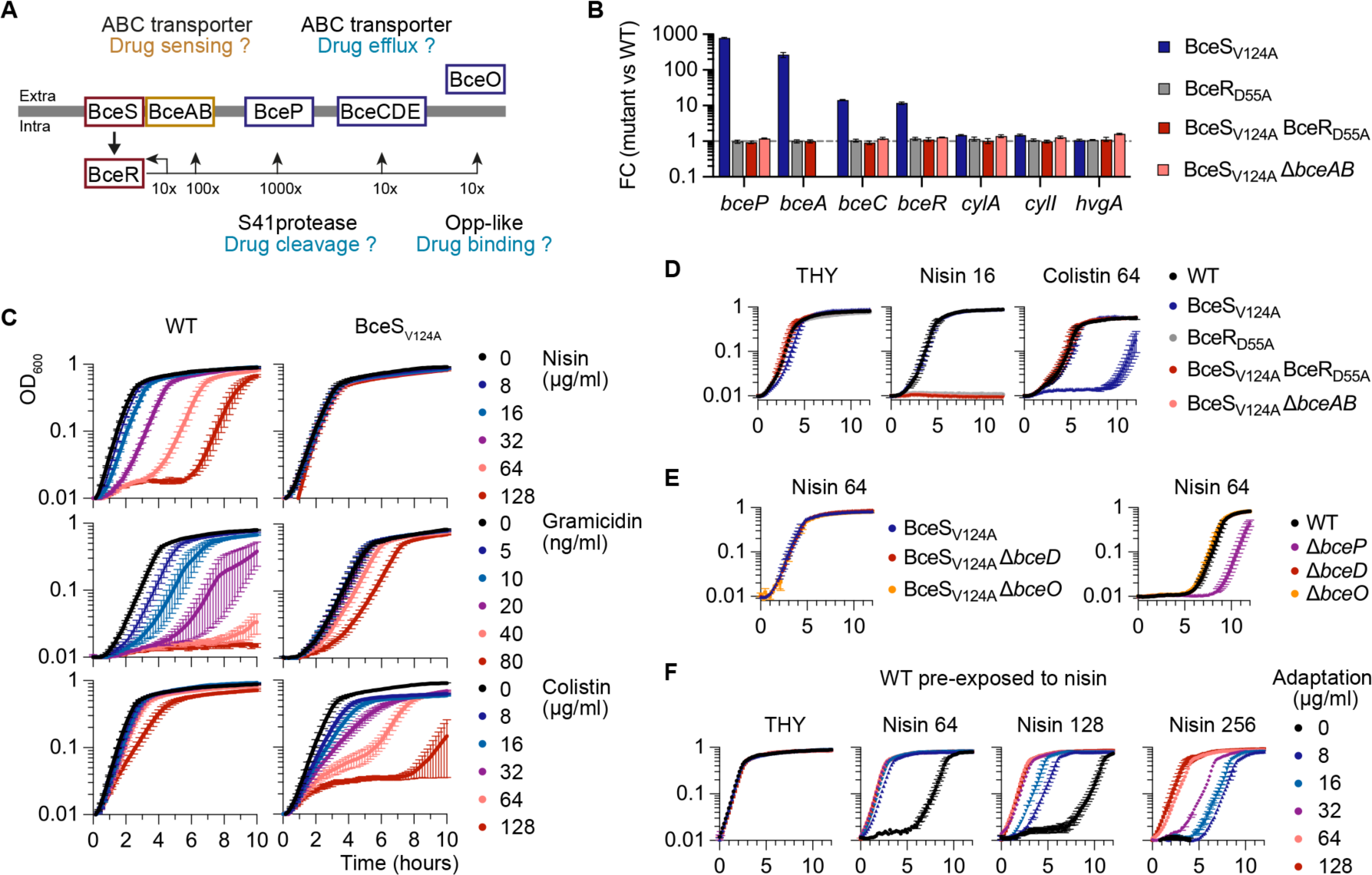
The BceRS three-component system controls an adaptive response. A. Schematic of the BceRS signaling pathway. The RNA-seq fold change scale in the BceS_V124A_ mutant is shown below the horizontal line. The BceAB transporter (yellow) is the third component of the regulatory system, predicted to sense and transduce the signal to BceS. Functions currently assigned to each component are indicated by question marks. B. BceAB is necessary to activate BceRS signaling in the absence of drugs. Fold changes during exponential growth in rich media were quantified by RT-qPCR in the activated HK^+^ mutant (BceS_V124A_), in mutants with a non-phosphorylable variant of the cognate regulator in the WT (BceR_D55A_) or activated (BceS_V124A_ BceR_D55A_) backgrounds, and in a BceAB transporter mutant in the activated background (BceS_V124A_ Δ*bceAB*). Bars represent the mean and SD of biological replicate (n = 3). C. Growth curves of the WT and activated BceS_V124A_ mutant in presence of increasing concentration of drugs. The curves represent the mean and SEM of biological replicates (n = 4). D. Drug susceptibilities of double mutants abolishing BceRS activation in the BceS_V124A_ mutant. The curves represent the mean and SEM of biological replicates (n = 3). E. Drug susceptibilities of Δ*bceP*, Δ*bceD*, and Δ*bceO* mutants in the activated mutants (left panel) and/or the WT strain (right panel). The curves represent the mean and SEM of biological replicates (n = 3). F. Growth curves of the WT strain pre-exposed to nisin. Early exponential growing WT strain were exposed for 4 hours to nisin (Adapt 0 to 128 µg/ml) in THY at 37°C. After washing and OD_600_ normalization, each culture is inoculated in fresh rich media (THY) and with increasing concentration of nisin (Nisin 64, 128, and 256 µg/ml). The curves represent the mean and SEM of biological replicates (n = 2).

Typically, this TCS family confers resistance to antimicrobials targeting lipid II cell wall metabolites such as nisin or bacitracin. Genetic activation of the pathway renders the BceS_V124A_ mutant insensitive to nisin, which has a marked effect on the lag phase but not on the growth rate of the WT strain, and increases resistance to gramicidin and, to a lesser extent, bacitracin (Fig. 5C and Supplementary Fig. S5). Interestingly, the BceS_V124A_ mutant is also more susceptible to antimicrobial peptides (colistin and polymyxin D) compared to the WT strain, while equally susceptible as the WT to vancomycin (Fig. 5C and Supplementary Fig. S5). Inactivation of the pathway in the BceS_V124A_ background by additional BceR_D55A_ or Δ*bceAB* mutations leads to nisin hyper-susceptibility while restoring WT level of colistin susceptibility (Fig. 5D). Nisin hyper-susceptibility is also observed for the single BceR_D55A_ mutant (Fig. 5D), a phenotype not linked to down expression of BceRS regulated genes (Fig. 5B).

To test the current model of nisin resistance based on drug efflux and cleavage, we inactivated the BceCDE transporter, the BceO substrate-binding protein, and the BceP extracellular protease (Fig. 5A). Deletion of Δ*bceD* and Δ*bceO* in the WT or BceS_V124A_ backgrounds has no impact on the nisin phenotypes of the respective parental strains (Fig. 5E), excluding a major function in drug export or binding. In contrast, the Δ*bceP* mutant is slightly more susceptible to nisin compared to the WT parental strain (Fig. 5E). However, deletion of *bceP* in the BceS_V124A_ background was always associated with secondary mutations inactivating the whole signaling pathway, this in five independent mutants (Supplementary Table S2). This suggests that the BceP extracellular S41 protease ^55^ has a buffering role when the pathway is activated, rather than directly cleaving drugs through an atypical mechanism, as previously suggested ^56^.

To test if BceRS regulates an adaptive response rather than a resistance mechanism per se, we pre-incubated the WT strain with nisin for four hours. Prior exposure to the drug decreases the lag-phase in a dose-dependent manner upon subsequent exposure to higher nisin concentrations (Fig. 5F). For instance, adaptation with 8 µg/ml nisin, a WT sub-inhibitory concentration, confers a BceS_V124A_-like resistance against a subsequent 64 µg/ml nisin challenge (Fig. 5F). More generally, prior adaptation with a given nisin concentration increases by a 4-fold factor the inhibiting concentration.

To summarize, the BceRS system is active in the absence of drug, required the BceAB transporter, and the basal level of constitutive activity is necessary and sufficient to cope with the stresses caused by sub-inhibitory nisin concentrations. The BceRS response is adaptative, inducible as required, and confers protection against structurally unrelated drugs targeting lipid II intermediates. However, individual regulated genes do not provide drug resistance and the response is constrained by its cost against antimicrobial peptides. Overall, this suggests that the BceRS system actively monitors and adjust surface-exposed lipid II metabolites, rather than directly detoxifying drugs or drug-lipid II complexes. In line with a drug-independent physiological function, the hypo-pigmented/haemolytic phenotypes of the BceS_V124A_ mutant (Fig. 3C), independent of CovRS regulation of the *cyl* operon (Fig. 5B), suggest cell surface alterations that could hinder interactions between the polyene backbone of the ß-h/c toxin and membranes ^46,48,57^.

## DISCUSSION

This study establishes the HK^+^ approach as a method of choice for characterising TCS signaling, both for mapping regulatory networks and for individual systems. This study was made possible by the conserved mechanism of HK phosphatase activity originally proposed _31,32_, which allowed a single residue to be targeted to activate the corresponding signaling pathway. By systematically testing all HisKA and HisKA_3 systems in a bacterium, we show the broad potential of this approach to reveal specialised, connected, and global regulatory systems covering the functional diversity that has evolved from a simple two-component architecture.

Targeting the HK phosphatase catalytic residue has the advantage of leaving a quasi-native system. The gain-of-function is solely dependent on the HK mutation, with no change to the RR and preservation of the physiological feedback loops. A second major advantage is that it bypasses the need for environmental signals, which are often unknown or confounding when having wide effect on bacterial physiology. In this respect, the SaeRS system is a remarkable example. Previous studies demonstrated SaeR regulation of *pbsP* and *bvaP* during vaginal colonization, among transcriptomic perturbations affecting nearly 40% of the genome ^49^. However, the regulon remained elusive due to lack of activation *in vitro* ^49^. The HK^+^ approach resolves the signaling pathway by revealing a specialized, and CovR-connected, pathway. Comparison with the well characterized *Staphylococcus aureus* homologous system ^58^ highlights the evolutionary divergence between regulatory circuits, particularly for those regulating host-pathogen interactions which need to be studied in each species.

An originality is the mechanism linking the SaeRS and CovRS systems. Complex regulatory wiring can be selected to mount co-ordinated responses, primary trough transcriptional cascades (a TCS regulating transcription of a second TCS) or connectors (usually a TCS-regulated transmembrane protein modulating the activity of a second TCS) ^59^. The C-peptide of adhesins can act as connectors when the transmembrane end remaining after cleavage of the LPxTG motif by sortase A interacts with a histidine kinase ^53^. The mechanism differs in GBS in which the PbsP adhesin domain acts as an extracellular signaling molecule to activate CovR signaling, independently of cell wall anchoring. We hypothesize that the lysin- rich and positively charged PbsP adhesin interacts with CovS, with the co-regulatory proteins Abx1 and Stk1 ^47,60^, or with the negatively charged membrane, recalling the activation of the homologous CovRS system in *Streptococcus pyogenes* by cationic peptides ^61^. To complete the regulatory circuit, CovR has previously been shown to repress *pbsP* in a strain-specific manner _45,50,51_. The intertwining of SaeR and CovR signaling through PbsP constitutes an adaptive mechanism for balancing adhesion and invasion and could contribute to the phenotypic and pathogenicity variabilities observed within the species.

HK^+^ mutations resolve TCS regulatory networks but reveal discrepancies in the activation of signaling pathways. While primary sequence analysis of TCSs did not uncover specific motifs correlating with high, intermediate, or low pathway activation, two underlying factors may dampen the effect of HK^+^ mutations. First, HK kinase activity can be inhibited by interacting proteins, such as the small LiaF protein inhibiting LiaS ^62,63^ and the Pst/PhoU proteins inhibiting PhoRS ^64,65^. Genes encoding co-regulatory proteins are often themselves regulated by the TCS, creating feedback loops that lock HKs in kinase-deficient conformation and thus obliterate the effect of HK^+^ mutations. Second, intermediary activation of the RelRS and CiaRH pathways suggests buffering mechanisms for TCSs regulating multiple independent loci and integrated cellular response. However, detailed analysis is required to decipher phosphorylation dynamics in each phosphatase-deficient HK^+^ and correlate *in vivo* RR phosphorylation with regulatory network activation, considering variable factors like the source of RR phosphorylation (kinase activity of the HK^+^ variant, cross-talk by other HK, small metabolites) and specific spontaneous dephosphorylation rates ^27,66,67^.

The systematic approach validates the conservation of the dephosphorylation mechanism. It also uncovers an unanticipated activation of the BceRS system with a degenerate QMKV motif. Recent structural insights from *Bacillus subtilis* complexes into the membrane environment supports a highly dynamic model of interactions between the BceAB transporter and the BceS kinase- and phosphatase-competent conformations ^68,69^. Our results with the HK^+^ BceS indeed suggest that BceAB is necessary to stabilize the kinase-competent conformation of BceS. Alternatively, BceAB could also provide the catalytic residue on the models suggested for the auxiliary phosphatases RapH and SpoOE ^31,70^. At the phenotypic level, our results point towards a need-based mechanism of target protection, as recently suggested for Bce-like system _71-73_, and not towards a drug cleavage-exclusion mechanism as initially suggested ^74,75^. The target protection mechanism relies on the binding of lipid II intermediates on a binding pocket of BceAB ^68^. However, it is still unclear how the system releases lipid II when it is complexed with drugs. Our results suggest an alternative scenario in which BceAB constantly monitor free lipid II intermediate to minimize target exposure ^76,77^. This alternative is supported by the steady-state activity of the BceRS pathway in the absence of drugs and is compatible with a need-based mechanism. Further studies should test the entire BceRS pathway without relying on a lipid II-drug detoxification mechanism, but rather on a mechanism that maintain the steady-state level of free lipid II in presence of drugs.

To conclude, genetic activation by HK^+^ is a powerful approach to characterize positive regulation by TCS. It circumvents the major drawback of studying systems that are usually non activated in standard condition. Previous studies on individual TCSs have demonstrated the potential of the approach, but it was unfortunately not widely adopted to date. Our systematic analysis based on the conserved mechanism of phosphatase activity provides a blueprint to decipher signaling, response dynamic, evolution of gene regulation, and regulatory networks. The HK^+^ approach is recommended for the study of TCS in any species, either as a complement or as a first choice alongside classical deletion mutants.

## MATERIAL AND METHOD

### Strain, mutants and genome sequencing

The BM110 strain is a clinical isolate representative of the hypervirulent CC-17 clonal complex responsible of most neonatal meningitis ^78^. The 2.2 Mb annotated genome is available under the NCBI RefSeq reference NZ_LT714196. The standard growth condition is in Todd-Hewitt medium supplemented with 1% yeast extract and 50 mM Hepes pH7.4 (THY) incubated in static condition at 37°C.

Oligonucleotides and construction of vectors for site-directed mutagenesis and deletion are detailed in Supplementary Table S5 and S6, respectively. Splicing-by-overlap PCR with high-fidelity polymerase (Thermo Scientific Phusion Plus) were done with complementary primers containing the desired mutations. The final PCR products contain mutations (SNP or deletion) flanked on either side by 500 bp of sequence homologous to the targeted loci. Cloning is done by Gibson assembly in the pG1 thermosensitive shuttle vector, before transformation in *E. coli* XL1-blue (Stratagene) with erythromycin selection (150 µg/ml) at 37°C. Inserts are validated by Sanger sequencing (Eurofins Genomics).

Mutant construction in GBS were done by a three-steps process involving: 1) electroporation and selection of GBS transformants at 30°C with 5 µg/ml erythromycin (permissive replication of the vector); 2) chromosomal integration on THY at 37°C with 5 µg/ml erythromycin (non-permissive replication temperature); 3) de-recombination by serial passage in THY at 30°c without antibiotic follow by colonies picking on THY at 37°C. Erythromycin-susceptible colonies having lost the vector were tested by discriminatory PCR (MyTaq HS - Bioline) with specific oligonucleotides (Supplementary Table S5) to select mutant over WT genotypes.

Genomic DNA of at least two independent mutants for each construction were purified from 1 ml of culture following manufacturer instruction for Gram-positive bacteria (DNeasy Blood and Tissue – Qiagen) and sequenced (Illumina sequencing at Core facility or Eurofins Genomics). High quality reads in Fastq were mapped against the BM110 genomes (55-419 x coverage, mean 181 x) and analysed with Geneious Prime (2019.2.3 - Biomatters Ltd). Results of genome sequencing for all mutants used in this study are summarized in Supplementary Table S2.

### RNA sequencing

RNA purification, sequencing and analysis were conducted essentially as described for the characterization of the virulence regulator CovR ^45^. The 14 HK^+^ mutants have been split into two series of 8 strains (7 mutants and one WT strain) and RNA was purified using three independent replicate that were grown on different days. Overnight cultures were used to inoculate THY (1/50), and 10 ml of culture are harvested in exponential growth phase (OD_600_ = 0.5) after incubation at 37°C. Bacterial pellets are washed with cold PBS containing RNA stabilization reagents (RNAprotect, Qiagen) before flash freezing and storage at -80°C. Total RNA are extracted after cell wall mechanical lysis with 0.1 µm beads (Precellys Evolution, Bertin Technologies) in RNApro reagent (MP Biomedicals), and purified by chloroform extraction and ethanol precipitation.

Samples were treated to remove residual DNA (TURBO DNase, Ambion) before fluorescent-based quantification (Qubit RNA HS, Invitrogen) and quality validation (Agilent Bioanalyzer 2100). Depletion of rRNA (FastSelect Bacterial, Qiagen), libraries construction and sequencing were done following manufacturer instructions (TruSeq Stranded mRNA, NextSeq 500, Illumina). Single-end strand-specific 75 bp reads were cleaned (cutadapt v2.10) and mapped on the BM110 genome (Bowtie v2.5.1, with default parameters). Gene counts (featureCounts, v2.0.0, parameters: -t gene -g locus_tag -s 1) were analysed with R v4.0.5 and the Bioconductor package DESeq2 v1.30.1 ^79^. Normalization, dispersion, and statistical tests for differential expression were performed with independent filtering. For each comparison, raw p-values were adjusted using Benjamini and Hochberg multiple tests ^80^ and adjusted p-value lower than 0.005 were considered significant. Raw sequencing reads and statistical analysis are publicly available (GEO accession number GSE261394).

In addition to HK^+^ RNA-sequencing, we have included an independent CovS_T282A_ transcriptome that was done simultaneously with the CovR_D53A_ transcriptome ^45^, the latter being already reported altogether with CovR ChIP-seq experiment (GEO accession number GSE158049). Gene networks are represented with the open-source software Cytoscape ^81^.

### RT-qPCR and promoter activity

For validation, independent RNA purifications from biological triplicates were done using the same protocol, except that the cultures were grown on the same day and only 1 ml was harvested. Reverse transcription and quantitative PCR (iScript Reverse Transcription and SsoAdvanced Universal SYBR Green, BioRad) were done using specific primers (Supplementary Table S5). Fold changes are calculated for each target relative to the WT strain whose RNA were purified in parallel.

For promoter activities, promoters were amplified and cloned in the pTCV-lac vector containing a ß-galactosidase reporter (Supplementary Tables S4 and S5) and introduced in GBS. Reporter activity was quantified in microplate format by colorimetric assay with ONPG as substrate and permeabilized overnight cultures ^47^. Reaction kinetics at 28°C were followed by OD at 420 nm every 5 min (Tecan Infinite). Linear slopes (OD/min) were used to infer enzymatic activities and were normalized for the initial cell density (OD 600 nm) of each replicate.

### Growth curves and antibiotic susceptibilities

Growth curves are done in a volume of 150 µl of THY inoculated with diluted overnight cultures (1/500) in 96-wells microplates and incubated at 37°C with automatic recording of OD 600 nm every 10 minutes and 1 minute agitation by cycle (TECAN Infinite). Doubling times are determined by fitting non-linear regression with a Malthusian growth model (GraphPad Prism 10) in exponential phase (R^2^ > 0.99) for each replicate. Fitness is calculated by dividing the individual doubling time against the mean WT doubling time. For antibiotic susceptibilities, concentrated drugs (10 x) were added to an aliquot of the starting cultures and serial two-fold dilution were done in starting culture without drugs before incubation in the microplate reader.

Minimal Inhibitory Concentration (MIC) is done following EUCAST guidelines in Muller-Hinton Fastidious culture media (MH-F, Becton Dickinson) media using custom AST Sensititre 96 wells plates (ThermoScientific) and 18h of incubation at 37°C.

### ß-haemolytic activity

Columbia agar supplemented with 5% horse blood and Granada medium (BioMerieux) were used to visualize ß-haemolytic activity and pigmentation, respectively. Serial 10x dilutions of cultures were spotted on media and incubated in aerobic (Columbia) or anaerobic (Granada with AnaeroGen, Oxoid) conditions at 37°C. To highlight halo of lysis around colonies, images are converted to gray scale and processed (Photoshop, Adobe) to uniformly adjust contrast and brightness. Haemolytic titers were determined by a semi-quantitative method ^47^. Serial 2-fold dilution of cultures initially adjusted to 10^9^ CFU/ml in PBS were added (V/V) to 1% defibrinated horse blood (Oxoid) in PBS supplemented with 0.2% glucose. After 1 hour of incubation at 37°C, cells were gently pelleted and hemoglobin in supernatants quantified by optical absorbance at 420 nm. Haemolytic activity of each strain was defined as the minimum dilution that lysed at least 50% of red blood cells. Haemolytic titers are the ratio between the haemolytic activity of each replicate against the haemolytic activity of the WT strain. Haemolytic titers are then normalized against the WT strain (normalized WT titer = 1).

### RR phosphorylation level

Genes encoding RR were amplified and cloned by Gibson assembly (Supplementary Tables S4 and S5) in a custom-made pEX-CterFLAG vector containing a synthetic cassette with a translational initiation site, a flexible Gly-Ala linker, a 3xFLAG epitope, and a transcriptional terminator. Cassettes with genes of interest cloned in frame with the linker were excised with restriction enzymes and cloned into the anhydro-tetracycline (aTc) inducible expression vector pTCV_P_tetO 45_. Expression vectors were introduced in the corresponding HK^+^ mutants by electroporation with kanamycin selection. Total protein extracts were prepared from 45 ml of cultures in exponential phase in presence of 100 ng/ml aTc (Sigma) by mechanical lysis of bacterial pellet (Precellys Evolution) resuspend in cold TBS buffer with EDTA-free protease inhibitors (cOmplete, Roche). Following clearance by centrifugation, 15 µg of proteins were deposit in 12.5 % Phos-Tag SDS polyacrylamide gels (SuperSep Phos-Tag, Wako Pure Chemical Industries Ltd) in loading dye buffer without EDTA and without sample heating to avoid dephosphorylation of the labile aspartate ^82^. Electrophoresis (2 hours, 100V, 30 mA) in Tris-glycine buffer was performed on ice bath. Semi-dry transfer on nitrocellulose membrane (15 minutes, 15 V, Mini-Protean, BioRad) was follow by blocking (TBS buffer with 0.05% Tween20 and 5% BSA), and hybridization with rabbit polyclonal anti-FLAG antibodies (1:1500, Sigma F7425) and finally with secondary antibodies coupled to infrared dyes (1:15000, Li-Cor IRDye 800CW). After final washing in TBS buffer without Tween20, fluorescent signals were acquired (Odyssey Imager, Li-Cor). Ratio of phosphorylated and non- phosphorylated protein were analysed with ImageJ from three independent protein extracts.

### Data availability

Raw sequencing reads and statistical analysis have been deposited in the Gene Expression Omnibus (https://www.ncbi.nlm.nih.gov/geo/) under GEO accession number GSE261394.

## Acknowledgements

This study was supported by Agence Nationale de la Recherche (VirEvol - ANR-22-CE15- 0024), and the National Laboratory of Excellence program - Integrative Biology of Emerging Infectious Diseases (LabEx IBEID, ANR-10-LABX-62-IBEID). CC and MVM are recipients of National PhD grants from Ecole Doctorale BioSpc (ED562) - Université Paris Cité.

## SUPPLEMENTARY TABLES LEGENDS

Supplementary Table S1: TCS-encoding genes in the hypervirulent BM110 strain.

Supplementary Table S2: List of strains with sequenced genome.

Supplementary Table 3: Antibiotic susceptibility.

Supplementary Tables S4: Transcriptome analysis of the HK+ collection. S4A: All results - RNA-seq normalized count - Mean of biological triplicate. S4B: All results - Statistical analysis.

S4C: Genes excluded from the analysis (rRNA, transposase, mobile genetic elements, no expression).

S4D: Differentially expressed genes (DEG) - threshold p-adj < 0.05 S4E: Activated genes in the HK+ collection (FC > 3, p-adj < 0.0001) S4F: Repressed genes in the HK+ collection (FC < -3, p-adj < 0.0001)

S4G: Activated regulon in the HK+ collection (-3 > FC > 3, p-adj < 0.0001)

Supplementary Table S5: Oligonucleotides.

Supplementary Table S6: Vector and mutant construction.

---

**Supplementary Figure S1:**
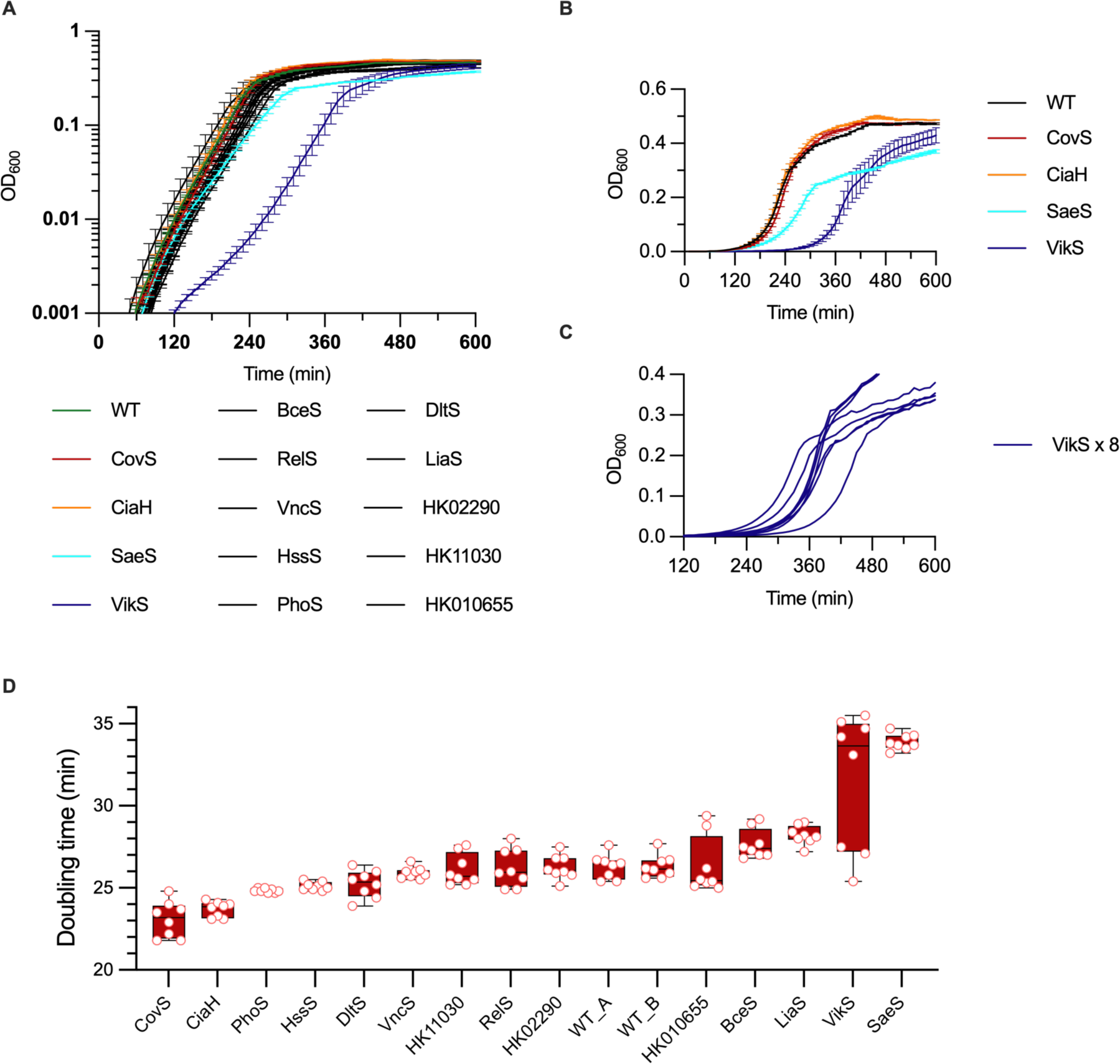
Growth phenotype of the HK^+^ collection. A. Growth curves of the HK^+^ collection. Cultures were done in rich media (THY) at 37°C in microplate starting from independent isolated colonies. Data represents the mean with SEM for each mutant (n = 8). B. Same data as in panel A in non-logarithmic scale highlighting the four mutants with significant phenotype. C. Individual growth curves of each replicate of the VikS_T221A_ mutants (n = 8). D. Doubling time in exponential growth phase. Malthusian non-linear fitting (r^2^ > 0.99) between OD_600_ 0.001 and 0.25 (panel A) were used to infer doubling time. Boxes represent the inter-quartile distance with median (horizontal lines), and the whiskers highlight minimal and maximal value (n = 8).

**Supplementary Figure S2:**
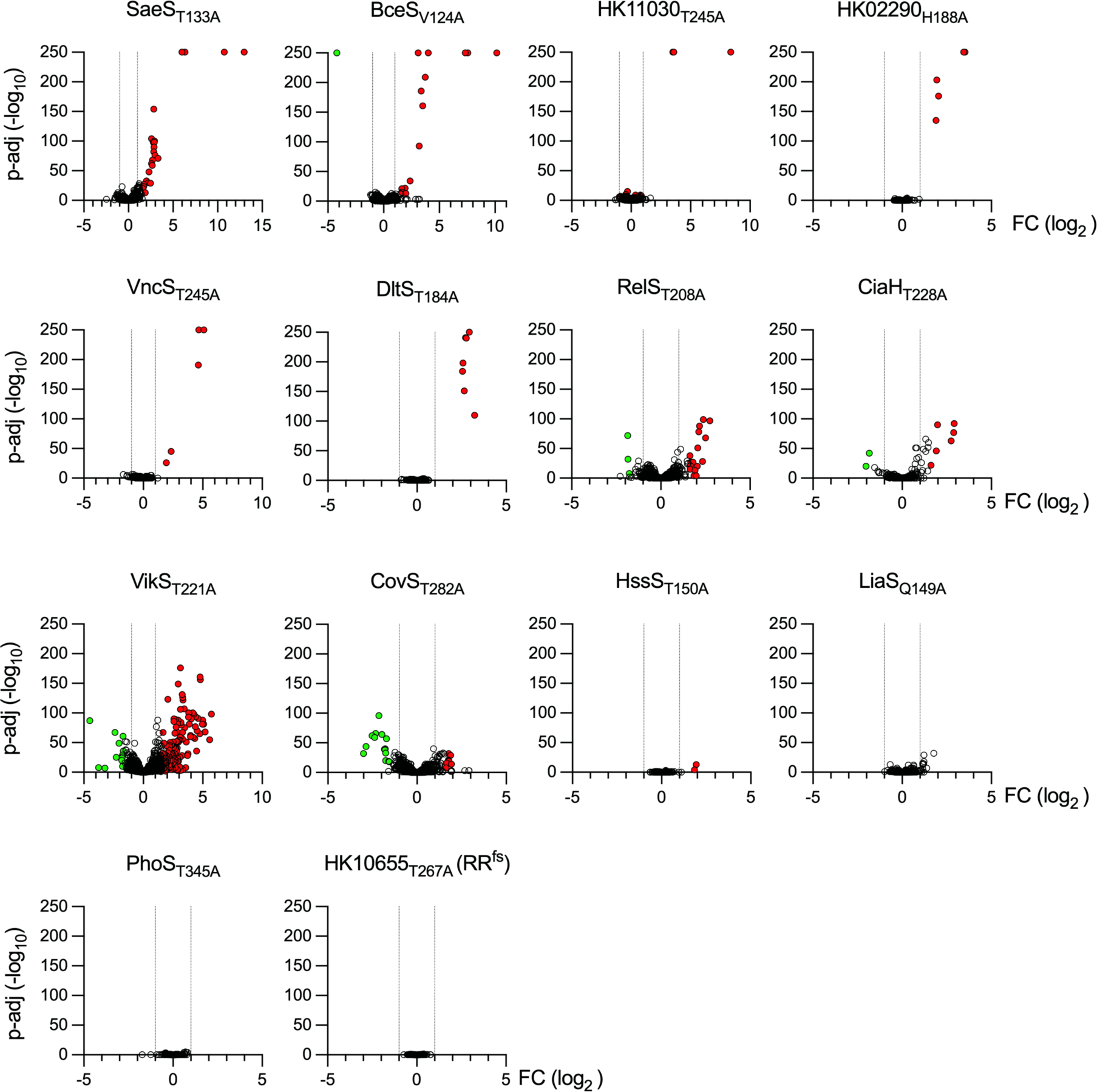
Significant differential gene expression in the HK^+^ collection Volcano plot of significant differential gene expression by RNA-seq in exponential growth phase at 37°C in rich media for each HK^+^ mutant against the WT strain. Coloured dots represent significantly activated (red) and repressed (green) genes (|FC| > 3, p-adj < 10^-4^), respectively.

**Supplementary Figure S3:**
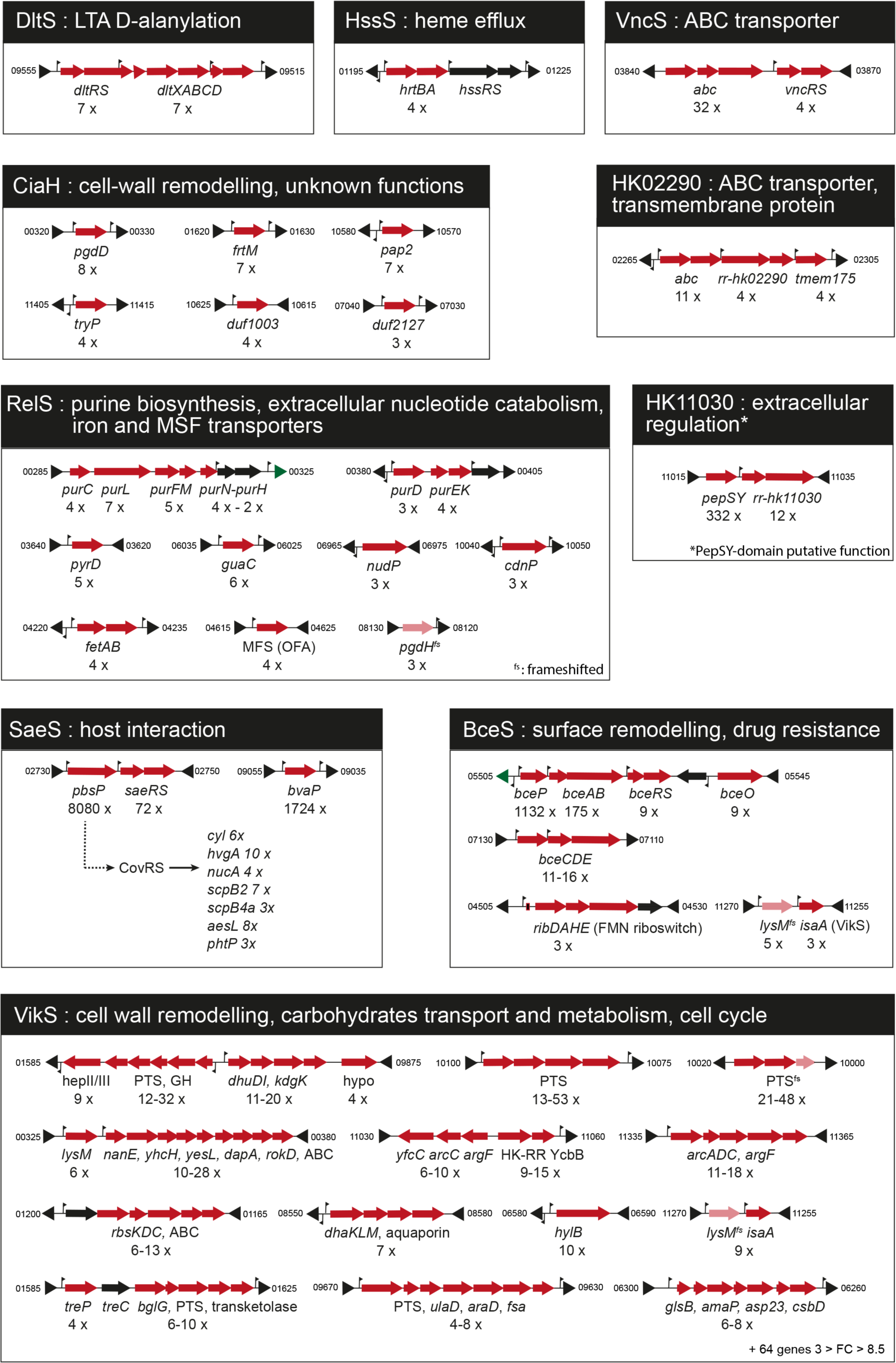
Activated chromosomal loci by HK^+^. Fold changes determined by RNA-seq are indicated below the activated genes (red arrows). Transcriptional start sites identified by genome wide TSS mapping are represented by vertical flags. NCBI genes ID bordering the loci are shown in a shortened form (e.g.: 11015 = BQ8897_RS11015). Frameshifted genes in the WT are marked (^fs^).

**Supplementary Figure S4:**
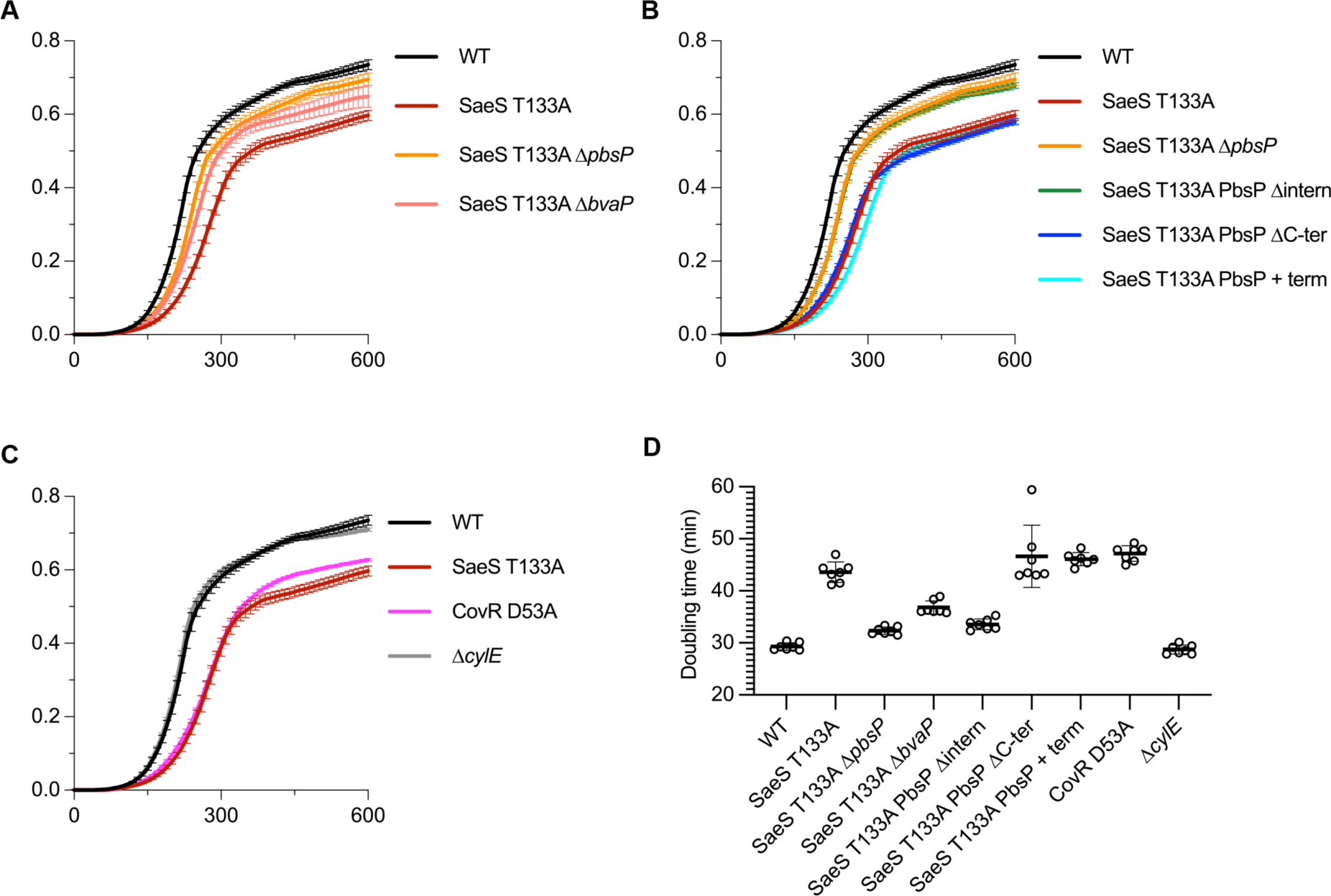
The fitness defect in SaeS_T133A_ is caused by the adhesin PbsP. A. Growth curves of the WT (black), SaeS_T133A_ activated mutant (red), and the double SaeS_T133A_ Δ*pbsP* (orange) and SaeS_T133A_ Δ*bvaP* (salmon) mutants. Data represent the mean and SD of a single experiment with pre-cultures inoculated with independent isolated colonies (n = 8). B. Same experiment with SaeS_T133A_ PbsP Δintern (green), SaeS_T133A_ PbsP ΔC-ter (blue), and SaeS_T133A_ + *pbsP* term (cyan). C. Same experiment with CovR_D53A_ (pink) and Δ*cylE* (grey). D. Corresponding doubling time in exponential growth phase. Malthusian non-linear fitting (r^2^ > 0.99) between OD_600_ 0.02 and 0.4 were used to infer doubling time. Dots represent biological replicate (n = 8) with mean and SD.

**Supplementary Figure S5:**
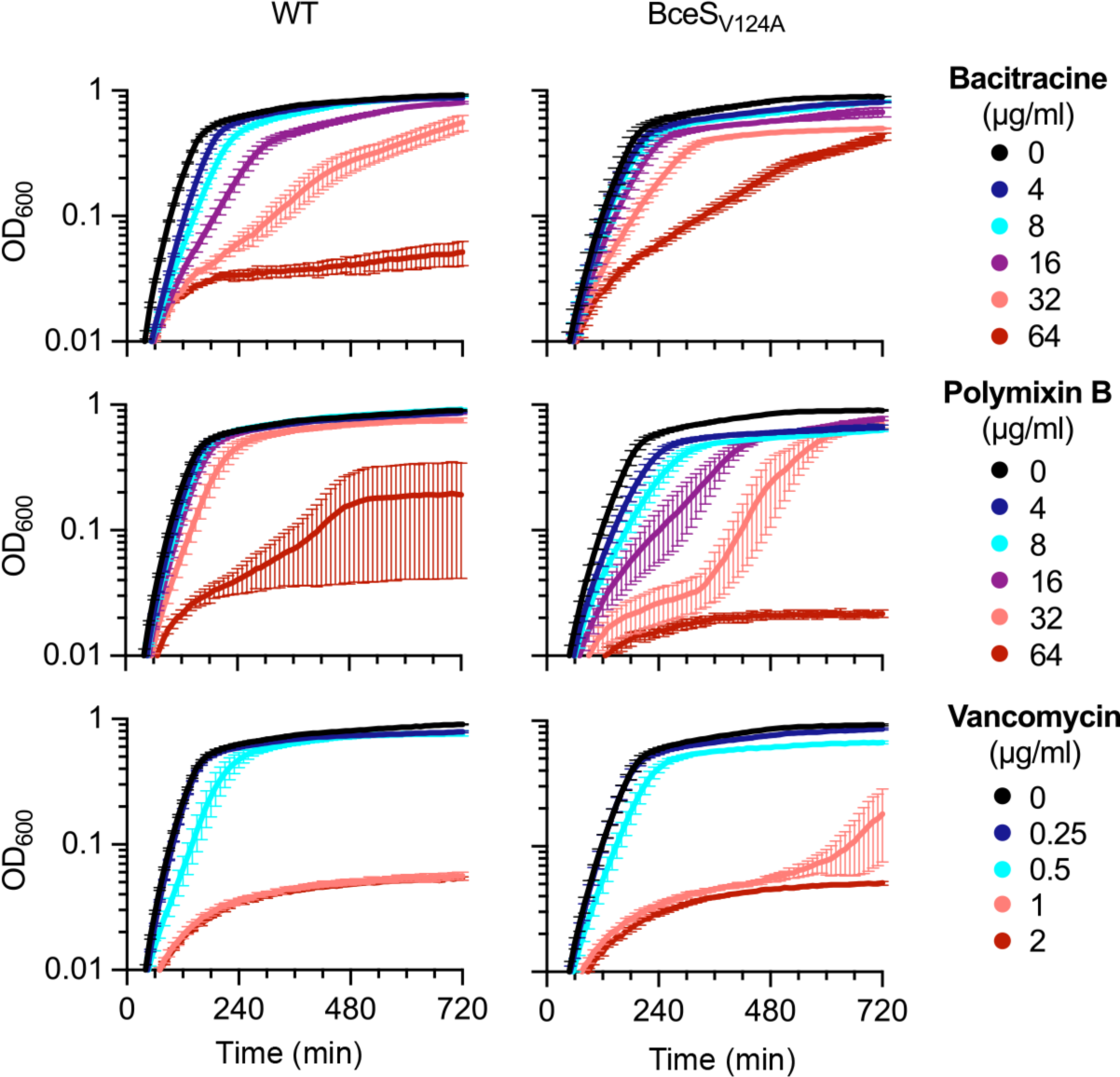
Drug susceptibilities of BceS_V124A_. Growth curves of the WT and activated BceS_V124A_ mutant in presence of increasing concentration of bacitracin, polymyxin B, or vancomycin. Curves are mean and SEM from biological replicate (n = 3).

